# Coordinated evolution between N2 neuraminidase and H1 and H3 hemagglutinin genes increased influenza A virus genetic diversity in swine

**DOI:** 10.1101/2020.05.29.123828

**Authors:** Michael A. Zeller, Jennifer Chang, Amy L. Vincent, Phillip C. Gauger, Tavis K. Anderson

**Author notes:** Address correspondence to: Tavis K. Anderson.

## Abstract

The neuraminidase (NA) and hemagglutinin (HA) of influenza A virus (IAV) are essential surface glycoproteins. In this study, the evolution of subtype N2 NA paired with H1 and H3 subtype HA in swine was evaluated to understand if genetic diversity of HA and NA were linked. Using time-scaled Bayesian phylodynamic analyses, the relationships of paired swine N2 with H1 or H3 from 2009 to 2018 were evaluated. These data demonstrated increased relative genetic diversity within the major N2 clades circulating in swine (N2.1998 between 2014-2017 and N2.2002 between 2010-2016). Relative genetic diversity of NA-HA pairs (e.g., N2.1998B/ H1.Delta1B) were correlated, suggesting intergene epistasis. Preferential pairing was observed among specific NA and HA genetic clades and this was associated with gene reassortment between cocirculating influenza A strains. Using the phylogenetic topology of inferred N2 trees, the expansion of genetic diversity in the NA gene was quantified and increases in diversity were observed subsequent to NA-HA reassortment events. The rate of evolution among NA-N2 clades and HA-H1 and HA-H3 clades were similar. The frequent regional movement of pigs and their influenza viruses is a possible explanation driving this pattern of drift, reassortment, and rapid evolution. Bayesian phylodynamic analyses demonstrated strong spatial patterns in N2 genetic diversity, and that frequent interstate movement of N2 clades homogenized diversity. The reassortment and evolution of NA and its influence on HA evolution may affect antigenic drift, impacting vaccine control programs and animal health.

## Introduction

Influenza A virus (IAV) is an important swine respiratory pathogen with high economic consequences in commercial swine production systems, with a risk for zoonotic transmission (Vincent, Ma, et al. 2008; Dykhuis-Haden, et al. 2012). Clinical signs of IAV in swine include cough, fever, lethargy, and a reduction of appetite that may lead to weight loss. IAV induced lung pathology often predisposes to secondary bacterial infections, resulting in further production losses and increased risk for mortality (Gramer 2006; Wang, et al. 2013). Due to the high morbidity associated with clinical disease, prevention and control of IAV is necessary to minimize animal suffering, mitigate production loss, and protect public health.

Vaccines have historically been formulated based on the hemagglutinin (HA) protein, which is the primary target of protective immune responses (Sandbulte, et al. 2015; Vincent, et al. 2017). Vaccines that contain HA antigens that are well-matched to the genetic diversity circulating IAV in swine have demonstrated efficacy, but this is greatly reduced when applied to control genetically drifted IAV (Vincent, Lager, et al. 2008). Further, the H1 and H3 HA subtypes of IAV that are detected in US swine contain at least 14 genetic clades with distinct antigenic properties (Anderson, et al. 2013; Zeller, Anderson, et al. 2018). Consequently, a potential strategy to increase the breadth of protection offered by IAV vaccines is the inclusion of additional proteins. Research has suggested that the neuraminidase (NA) gene may also contribute to vaccine efficacy through neuraminidase inhibiting antibodies (Monto and Kendal 1973; Sandbulte, et al. 2016).

N1 and N2 subtype NA genes circulate in the US, with the N2 subtype being detected in approximately two-thirds of swine IAV reported in surveillance programs (Anderson, et al. 2013; Zeller, Anderson, et al. 2018). Within these NA subtypes, there are two N1 genetic clades, including the classical swine lineage (N1.Classical) which emerged coincident with the 1918 H1N1 introduction into swine (Koen 1919), and a Eurasian swine lineage N1 gene (N1.Pandemic) that emerged in the US associated with the 2009 H1N1 human pandemic (Dawood 2009; Smith, et al. 2009). Contemporary N2 genetic clades circulating in swine are derived from human-to-swine transmission episodes: the first associated with human seasonal IAV in the late 1990s (N2.1998), and the second in the early 2000s (N2.2002). The N2.1998 lineage was introduced into swine with the triple reassortant IAV that evolved into the H3.Cluster IV HA (Zhou, et al. 1999). The N2.2002 lineage was introduced during a human to swine spillover coincident with the H1.Delta clades (Vincent, et al. 2009). Although veterinary diagnostic labs continue to occasionally detect sporadic human seasonal NA in swine (Zeller, Li, et al. 2018), these have yet to become widely established.

The quantification of genetic and antigenic diversity described in IAV in swine has largely focused on the HA gene. However, the NA and HA function in a coordinated effort to replicate within and transmit between hosts. Functionally, a balance in NA and HA cellular interactions are necessary for IAV infection to result in successful host-to-host transmission (Mitnaul, et al. 2000; Das, et al. 2011). Links between these genes have been documented in phylogenetic studies on human seasonal H3N2 which have suggested that mutation in one gene affects mutation in the other (i.e., inter-gene epistasis) (Neverov, et al. 2014). In swine, a similar type of evolutionary dynamic is plausible given the 14 different HA genetic clades and 4 NA lineages with considerable genetic diversity within each, but the degree to which this affects observed diversity is unknown.

The objective of this study was to assess the NA genetic diversity of IAV circulating in US swine and evaluate how evolution and reassortment to pair with new HA subtypes or clades may affect the genetic diversity and evolutionary trajectory of the N2 gene. Understanding how HA pairing and reassortment impacts the evolutionary dynamics and extent of genetic diversity of the NA in swine IAV is critical to understanding how the genomic context affects the emergence and transmission of IAV in swine and will facilitate control efforts that include NA in vaccine strain selection.

## Results

### Increased N2 relative genetic diversity

Effective population size (EPS) was used to quantify changes in relative genetic diversity in swine IAV N2 lineages and clades (Figure 1). The N2.1998 demonstrated an increase in diversity between 2014 and 2017 whereas the N2.2002 lineage demonstrated an increase in diversity between 2010 and 2016. Four statistically supported monophyletic clades were detected within these broad swine N2 lineages, N2.1998 split into N2.1998A and N2.1998B circa 2008 (2006.46 – 2008.78 95% HPD), and N2.2002 split into N2.2002A and N2.2002B circa 2006 (2004.92 – 2007.14 95% HPD). The relative genetic diversity for each of these clades are presented in Figure S1. The average within genetic clade distance was 3% and the between clade distance was greater than 6% for each of the monophyletic clades in each lineage; but, there was clade specific variation in diversity. The N2.1998A had a constant EPS, compared to the increasing EPS observed in the N2.1998B clade. The N2.2002A demonstrated an initial increase that peaked in mid-2012, before being gradually superseded by the N2.2002B by mid-2014 and peaking in 2015. These data suggest a general trend of increasing relative genetic diversity observed in both N2 lineages. The N2.1998B clade appears to be the major driver of the increased EPS of the N2.1998 lineage whereas the N2.2002A and N2.2002B demonstrated different temporal trends that both contributed to the overall increase in EPS across the study period.

**Figure 1.**
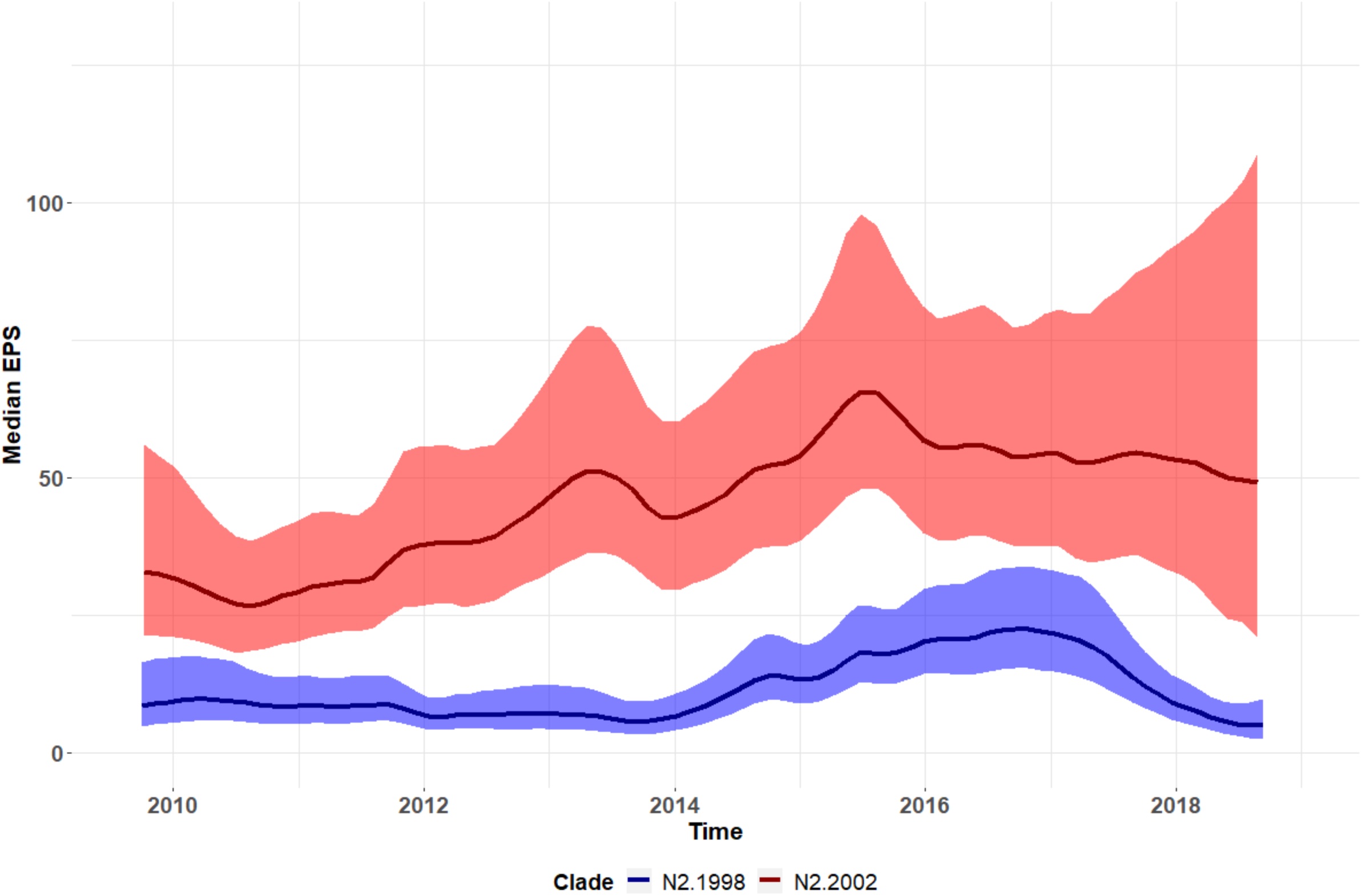
Relative genetic diversity, measured as effective population size (EPS), of the predominant N2 neuraminidase clades in influenza A viruses detected from 2009-2018 in swine in the United States (N2.1998 in blue and N2.2002 in red). Median EPS denoted by solid colored lines with the 95% HPD shaded in the same color. Relative genetic diversity increased linearly in the N2-1998 between 2014 and 2017, and between 2010 and 2016 in the N2.2002.

### N2 and HA reassortment and non-random pairing

NA-HA pairing was inferred from the N2 trees annotated by HA clade. Distinct clustering of HA clade groups was evident, along with shifts in HA clade annotation on the NA genes as a likely result of reassortment. We defined establishment of the IAV HA and N2 clade pairing within the swine population when they were observed with greater than 10 detections across multiple years. Nine such reassortment events were found and annotated (Figure 2). Longer branch lengths were observed after reassortment, suggesting an increase in N2 genetic diversity. A Chow test (Chow 1960) was used to statistically validate if there was a change in the rate of mutation of the N2 neuraminidase before and after HA and NA reassortment (Figure S2). Of the nine reassortment events observed, three had significant support for differences in evolutionary rate. The reassortment event labeled as “1” in Figure 2A, a transition in N2.1998A from H3.ClusterIVF to H1.Delta1B (p < 0.001) (Figure S2A), showed the rate of mutation slowed following reassortment. The reassortment event labeled as “7” in Figure 2B described an N2.2002B paired with H3.ClusterIVB reassorting to Cluster IVA (p < 0.001) (Figure S2E): in this case, the rate of mutation increased following reassortment. The reassortment event labeled “8” in Figure 2B was an N2.2002B gene that changed pairing from H3.ClusterIVA to H1.Delta1A (p < 0.001) (Figure S2F); for this event, the rate of mutation remained the same after reassortment, but there was a increase in branch length to the oldest gene in the genetic clade of these pairings. The remaining reassortment events with onward transmission in the US swine population shown in Figure 2 had a general trend towards increased mutation rates following reassortment but this was not statistically significant (Fig S2).

**Figure 2.**
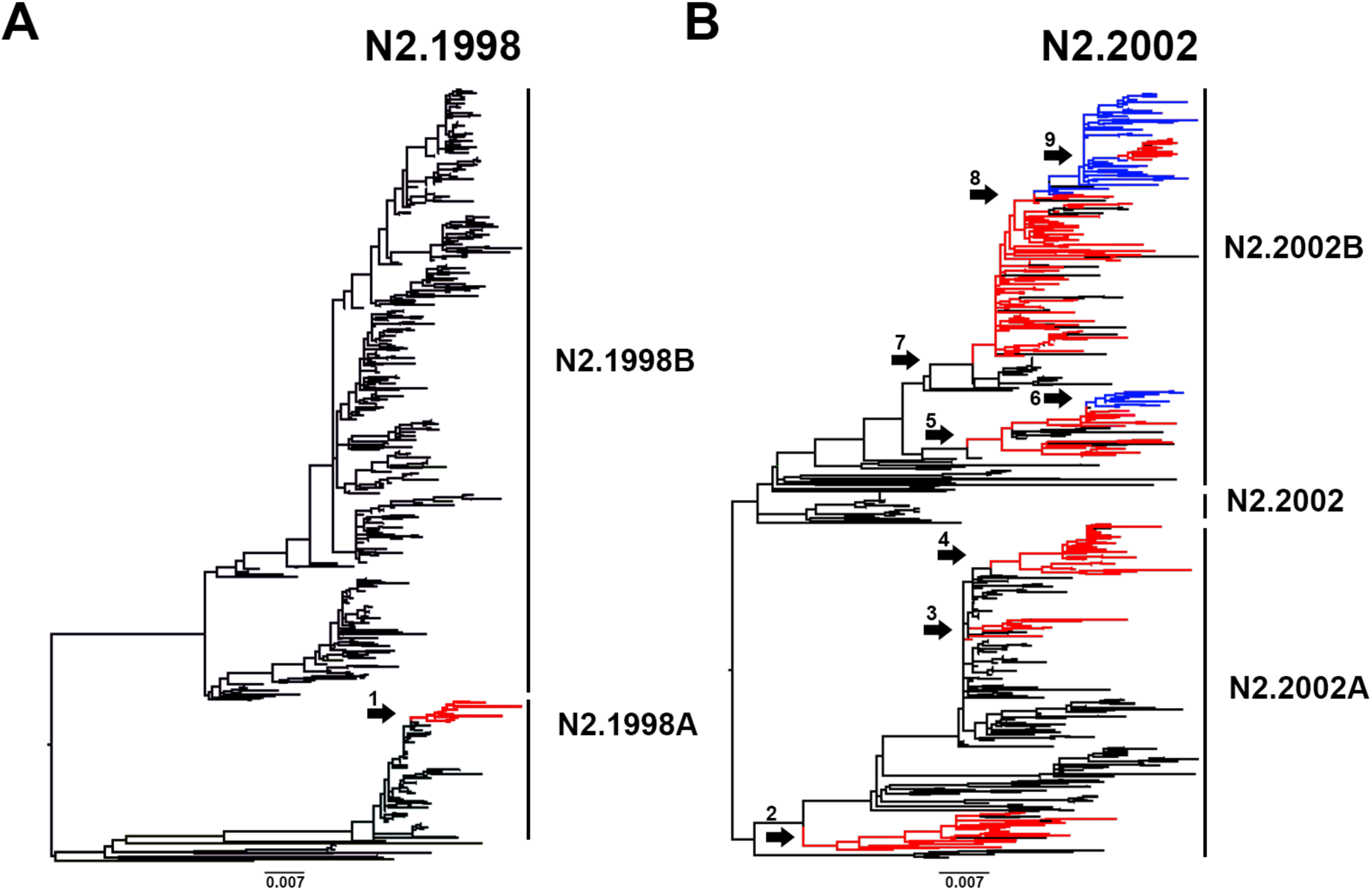
Maximum likelihood trees of the (A) N2.1998 and (B) N2.2002 lineage neuraminidase (NA) genes with N2 sequences from August 2009 to July 2018. Phylogenetic trees show sustained (> 10 detections) paired hemagglutinin (HA) clade transitions through colored changes, being either black to red or red to blue. NA genetic clades are annotated on the right of each panel. A numbered arrow indicates a reassortment event where an N2 gene was paired with a new HA genetic clade with subsequent sustained detection in the swine population. The 10 numbered arrows represent HA gene exchanges as follows: (1) H3 C-IVF to H1 Delta1B; (2) H1 Delta1B to H1 Delta1A; (3) H1 Delta1B to H3 C-IVA; (4) H1 Delta1B to H3 2010.1; (5) H3 C-IVB to H1 Alpha; (6) H1 Alpha to H3 2010.1; (7) H3 C-IVB to H3 C-IVA; (8) H3 C-IVA to H1 Delta1A; and (9) H1 Delta1A to H3 2010.1. Each sustained reassortment event demonstrated increased genetic divergence, visualized as longer branch lengths in the phylogeny. The scale bar represents approximately 10 nucleotide changes in the neuraminidase gene.

A tanglegram consisting of a subsampled pairing of N2 and HA sequences was generated. Lines connecting N2 to their constituent HA within strain tended to be paired by clade, suggesting these pairings were not random (Figure 3). In Figure 3, the connecting lines indicate NA-HA pairings: the frequency of detection and the topology of these trees demonstrated some pairing pattern shifts that subsequently became fixed in the population (Figure 3). Prior to 2013, detections of H1.Delta1A were paired with N2.2002A. Following reassortment in 2013, this HA gene was frequently paired with N2.2002B and became the predominant H1.Delta1A pair by 2015 (Figure S3A). Prior to 2011 H3.ClusterIVA were paired with N2.2002A, and after 2011 this HA was paired with N2.2002B and became the predominant pairing by 2013 (Figure S3B). The H3.2010.1 was paired with the N2.2002A predominantly from 2014-2017, after which N2.2002B became predominantly detected (Figure S3C).

**Figure 3.**
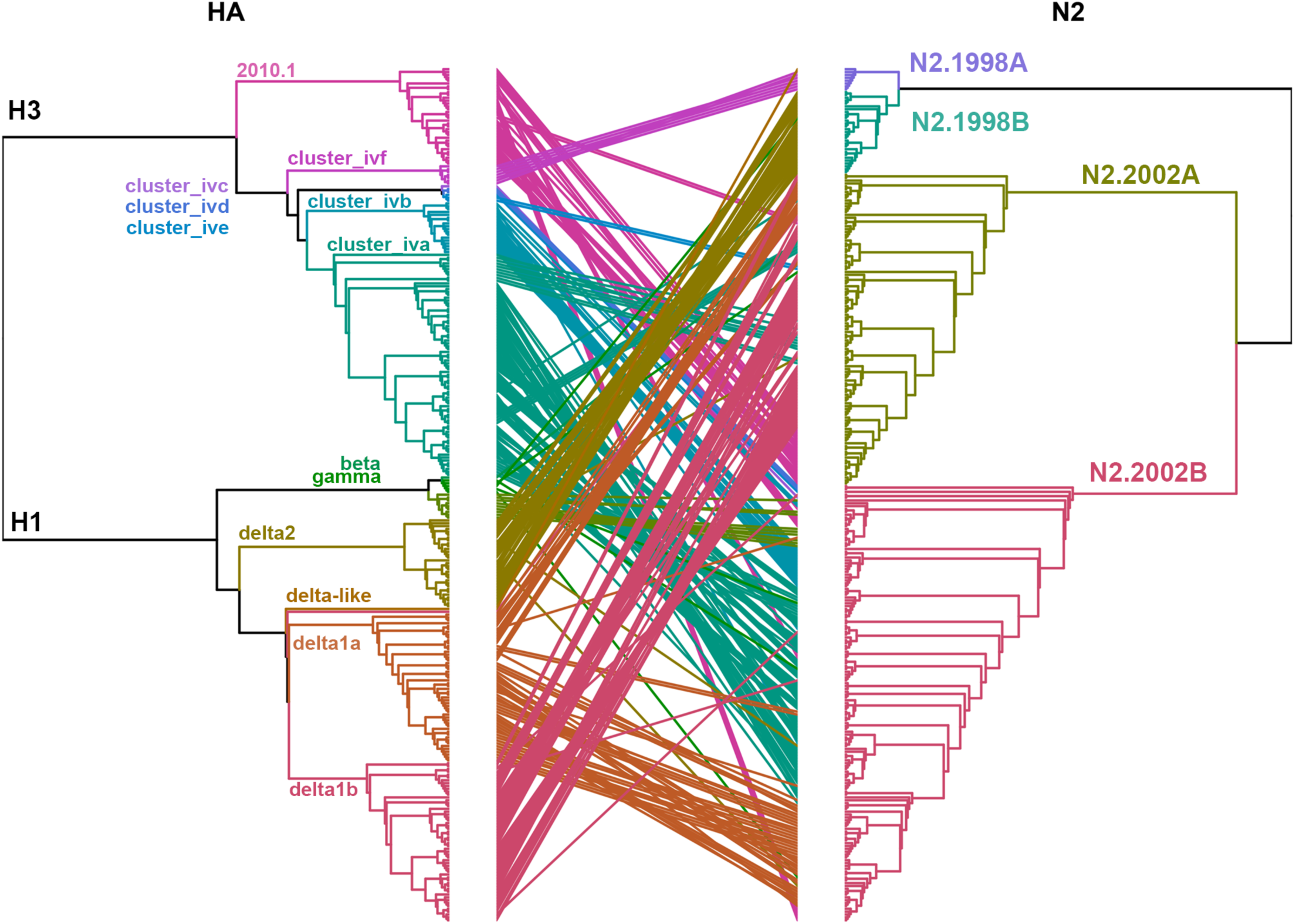
Tanglegram of the hemagglutinin (HA) and paired N2 neuraminidase (NA) genes in US swine influenza A virus. HA and N2 NA lineages are indicated by color and are presented on defining branches of the phylogeny. Connecting lines in the tanglegram are colored by the HA clade and demonstrate a link between HA and N2 present in the same virus. Maximum likelihood trees are presented as cladograms with branch lengths scaled proportionally to the changes per site. Multiple, repetitive connecting lines between specific HA and N2 clades was suggestive of non-random pairing. Reassortment is indicated when connecting lines move across multiple clades. For example, the H3.Cluster IVA and H1.Delta2 demonstrated connecting lines with multiple N2 clades, suggesting reassortment events occurred that subsequently became fixed in the swine population. Genetic divergence of HA was observed during these reassortment events based on branch length.

The non-random pairing observed in the tanglegram was statistically significant (chi-squared (p < 0.0001); Fisher’s exact test (p < 0.0001)). The distribution of N2-HA pairings (Figure S4A) were subjected to a post hoc analysis using the standardized residuals from the chi-squared test (Figure S4B). This demonstrated disproportionate pairing of N2 clades with their constituent HAs. N2.1998A were predominantly paired with H1.Delta1B and H3.ClusterIVF. This pairing demonstrated a temporal trend, as most of the N2.1998A detections were associated with the H1.Delta1B rather than H3.ClusterIVF following reassortment (Figure 2). N2.1998B were almost exclusively paired with H1.Delta2. The N2.2002A were disproportionately paired with H1.Delta1B or H3.ClusterIVE (Figure S4B): both clades were infrequently detected (Figure S4A). The N2.2002B was disproportionately paired with H1.Delta1A, H1.Alpha, and the H3.ClusterIVA-D (Figure S4B). The H1.Delta1A and H3.ClusterIVA HA clades were also associated with N2.2002A indicating these HA genes permit multiple N2 clade pairings (Figure S4B). The H3.2010.1 clade viruses were paired with either N2.2002A or N2.2002B in approximately equal numbers, but data collected over the last 2 years demonstrated an increase of N2.2002B pairing (Figure S3C). The non-random pairing observed between N2 and HA genes through the chi-squared test indicated a potential correlation in relative genetic diversity of each gene. The relative genetic diversity of the N2.1998B clade paired with the H1.Delta2 clade and the N2.2002A paired with the H1.Delta1 clade were selected for analysis due to their near exclusive pairing. Detection of the H1.Delta2 paired with the N2.1998B first occurred in 2011 and thereafter became the predominant N2-HA pairing of these clades. After 2012, the EPS of the N2.1998B and the H1.Delta2 became superimposed both temporally and in magnitude, suggesting that changes in the genetic diversity of these two genes are linked (Figure S5A). Paired N2.2002A and H1.Delta1B demonstrated similar temporal aspects and magnitude of their respective measures of relative genetic diversity (Figure S5B).

Similarity in the relative genetic diversity of N2-HA pairs indicated that the mutation rate of N2 clades and paired HA clades are likely similar. Comparing the mean substitution rate per codon of N2 clades and select HA clades demonstrated similar mutation rates, N2.1998A: 0.0044, N2.1998B: 0.0046, N2.2002A: 0.0041, N2.2002B: 0.0045, H1.Gamma: 0.0047, H1.Delta2: 0.0049, H3.ClusterIVA: 0.0049, H3.2010.1: 0.0050 sites/mutation/year (Figure S6). Collectively, the overall NA mutation rate was slightly lower than the overall HA mutation rate, with a difference of 0.000475 sites/mutation/year.

### Diversifying selection in the N2

A mixed effects model of evolution selection analysis demonstrated that multiple amino acid sites were under positive selection for each N2 clade (Table 1). The N2.1998A had 4 amino acid sites under positive selection in both the transmembrane and globular head regions. Although the analysis indicated position 414 as under positive selection, glycine was present at position 414 in all but three N2.1998.A genes and did not appear to be truly under positive selection. The N2.1998B had one transmembrane, five stalk, and four globular head sites under positive selection. There were two distinct monophyletic groups nested within the N2.1998B clade with the sites under positive selection also split across each clade, suggesting these two groups diverged prior to 2009 (the earliest data used in our analysis). The N2.2002A had one stalk and three globular head sites under positive selection, but no transmembrane sites. The N2.2002B had one transmembrane, one stalk, and seven globular head sites under positive selection. These data supporting the presence of positive selection across each of the N2 NA clades indicates that functional amino acid mutations are likely occurring in the gene as it is evolving.

**Table 1.**
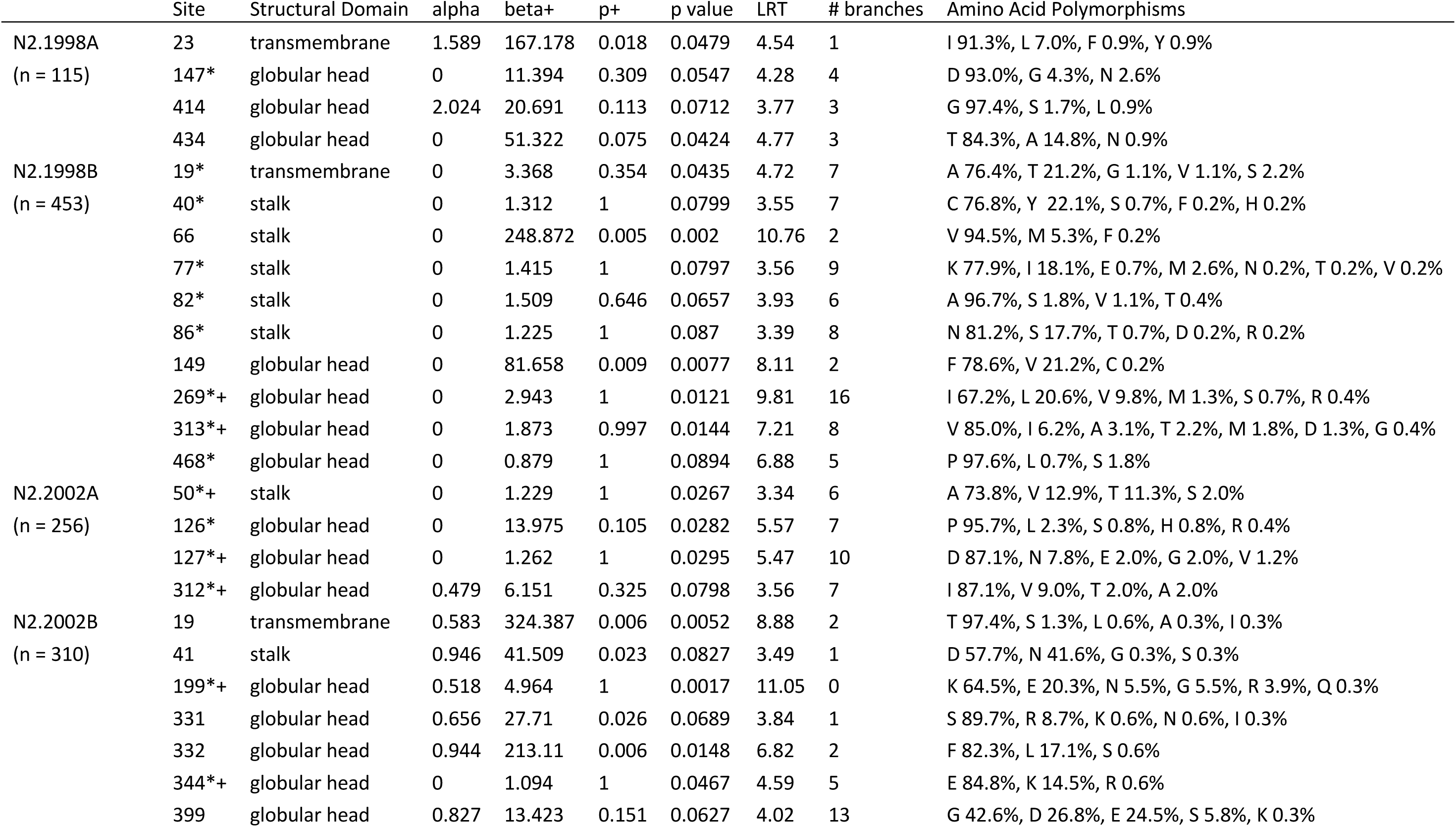
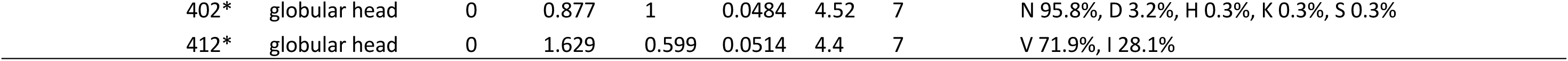
Amino acid sites detected as undergoing positive selection in the neuraminidase N2 gene of swine influenza A viruses. Instances of episodic and pervasive selection were determined using a mixed-effects model of evolution (MEME) part of the HyPhy package. Sites under positive selection were also assessed with the FEL and SLAC methods in HyPhy. Sites common between MEME and FEL were annotated with ‘*’ and those overlapping between MEM and SLAC were annotated with ‘+.’

### Directionality of reassortment

Reassortment resulting in HA and N2 gene pairings did not appear to be random (Figure 4). There was limited reassortment of the N2.1998A clade NA genes from H3.ClusterIVF to H1.Delta1B viruses (Figure 4). The N2.1998B clade genes were almost exclusively paired with H1.Delta2, with very few sporadic dead-end reassortment events detected. The few reassortment events within the N2.1998B clade were given high and uniform posterior probabilities as well as Bayes factors, but our maximum likelihood trees indicated that despite N2.1998B reassortment, it was not consistently detection or sustained in swine. The N2.2002A genes demonstrated reassortment from H1.Delta1A, H1.Delta1B, H3.ClusterIVA, and H3.2010.1 to multiple other HA clades, indicating the N2.2002A genes may pair with multiple HA genes. The N2.2002B demonstrated the greatest breadth of reassortment with the gene pairing with H1.Delta1A, H1.Alpha, H3.ClusterIVA, H3.ClusterIVB, and H3.2010.1. These results indicate that some NA genes are permissive of multiple HAs (i.e., approaching random), whereas other NA genes are only detected with specific HA genes. This Bayesian approach is consistent with the clade pairing patterns observed in the tanglegram and chi-squared analysis (Figure 3 and Figure S5).

**Figure 4.**
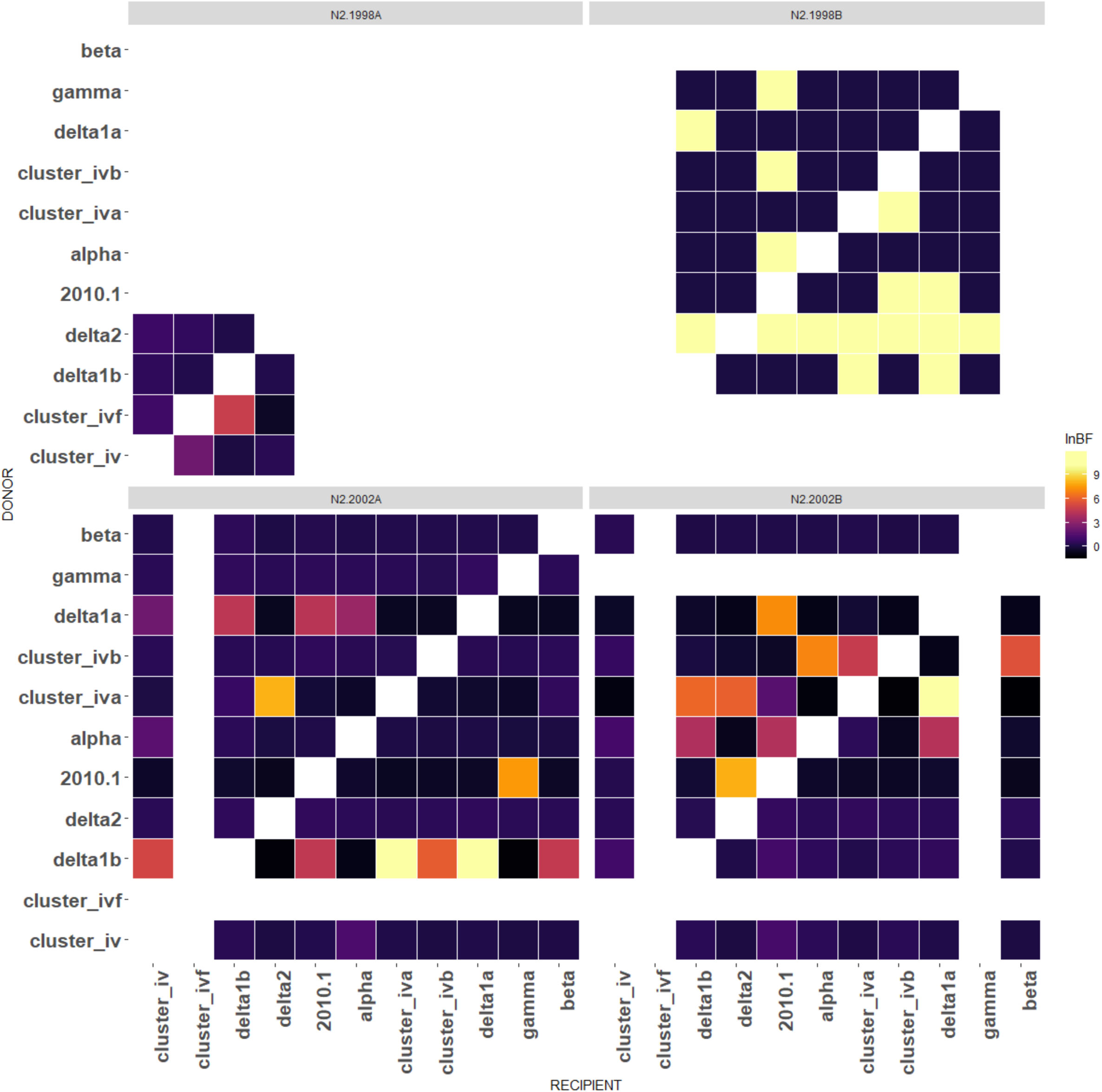
Bayes factors (Ln BF) representing the confidence of transition events of the hemagglutinin (HA) gene by clade for neuraminidase (NA) gene by N2 clades from 2009 to 2018. Higher Ln Bayes factors represent in warmer colors are higher evidence for reassortment; uncolored squares present an absence of the transition between genes. The N2.1998A clade demonstrated the least frequent reassortment of the N2 clades, but there was a transition from H3.ClusterIVF to H1.Delta1B. The small numbers of detected reassortment events for N2.1998B had strong support due to near exclusive pairing between N2.1998B and H1.Delta2. Both N2.2002A and N2.2002B NA genes demonstrated evidence for multiple reassortment events.

### Spatial dissemination of N2 genetic clades

The spatial detection of N2 clades from 2009-2018 indicated differences in geographic distribution (chi-squared test p < 0.0001: Figure 5A) that reflect general patterns of swine agriculture in the US. Viruses containing N2.1998A had the fewest detections and had limited geographic distribution in the Midwest, primarily in Iowa or adjacent states. The N2.1998B was detected most frequently in the Southeast (North Carolina), with a few detections in the Midwest including Indiana and Iowa. The N2.2002A was detected most frequently in the Midwest and was rarely detected in other regions of the US. The N2.2002B was detected in the Midwest as well as the Southeast (North Carolina). On a state basis, differences in detection of viruses containing specific NA clades was observed, for example viruses detected in Iowa contained NA clades in descending order of frequency: N2.2002B, N2.2002A, then N2.1998B.

**Figure 5.**
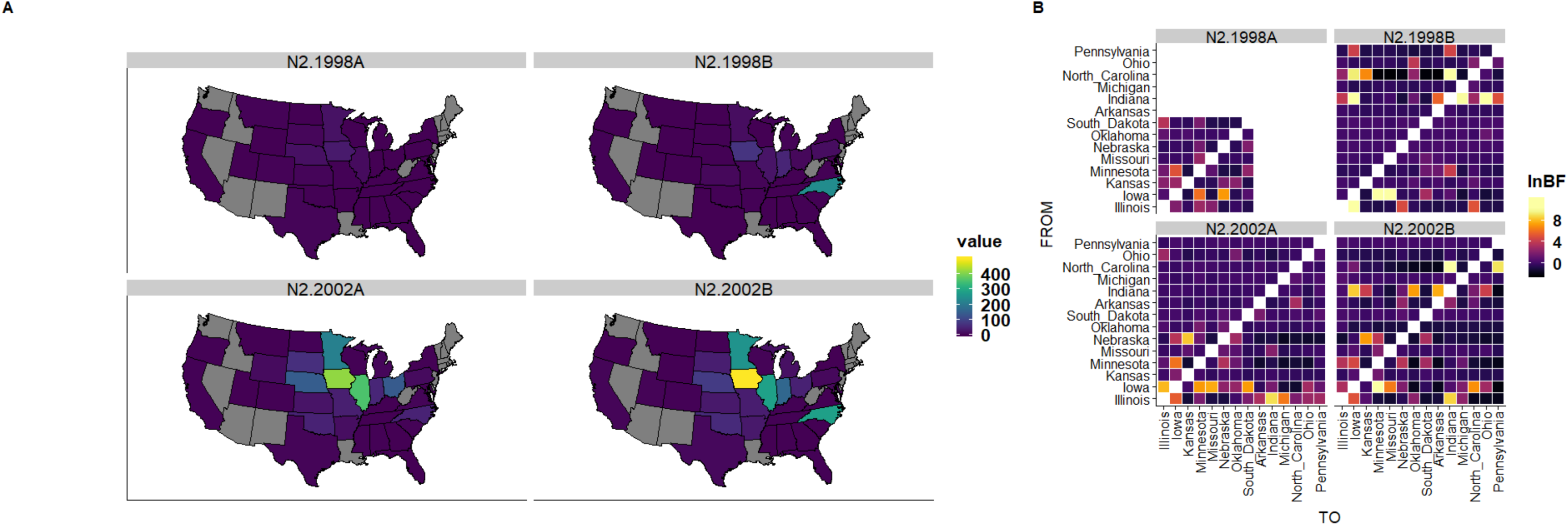
(A) Geographic representation of each N2 clade from 2009-2018. N2 clade was dependent on location (chi squared: p < 0.001). N2.1998A was observed with few detections in limited locations in the Midwest. N2.1998B is detected primarily in North Carolina, with few detections present in the Midwest including Indiana and Iowa. N2.2002A was detected in the Midwest but was rarely detected in North Carolina. N2.2002B was detected in the Midwest as well as North Carolina. (B) Bayes factors for inferred state-to-state movement of N2 clades demonstrating the frequency with which particular genetic clades are moving from particular states on the y-axis to other states on the x-axis.

Interstate N2 gene movement, inferred by Bayesian phylodynamic techniques (Figure 5B), demonstrated that even infrequently detected N2 had geographic movement despite the low overall distribution presented in Figure 5A. IAV containing N2.1998A moved between Minnesota, Iowa, and Nebraska. IAV with N2.1998B moved from North Carolina into the Midwest as well as from Iowa to Minnesota and Missouri. Despite movement to the Midwest, the N2.1998B was not frequently detected in that region. IAV with N2.2002A were moved (likely transported with live swine) from Iowa and Indiana to multiple locations throughout the Midwest, including Nebraska and Kansas. IAV with N2.2002B moved from North Carolina to Indiana and Pennsylvania, with Indiana, Iowa, and Illinois also acting as sources of N2 gene distribution. Nebraska appeared as a possible source for N2.2002B to move into Kansas. Despite the inference of transmission of almost all N2 genetic clades to all geographic regions across the US, the detection of N2 clades within surveillance data suggested that many of these clades do not persist.

## Discussion

Multiple cocirculating clades of swine IAV NA were reported, with a significant amount of genetic diversity that challenges control efforts (Anderson, et al. 2013; Zeller, Anderson, et al. 2018). Our data demonstrated that diversity in the NA gene has increased dramatically over the past 10 years. Previously described N2.1998 and N2.2002 lineages genetically diverged circa 2008 and 2006 respectively, warranting the naming of four statistically supported monophyletic clades (N2.1998A, N2.1998B, N2.2002A, N2.2002B). Our data suggested that this observed diversity has occurred due to the concurrent forces of drift, reassortment that resulted in novel NA-HA pairings, and interstate movement of IAV. These dynamics played a critical role in establishing predominant NA-HA pairings. The interstate movement of IAV with pigs may perpetuate the cycle of drift, reassortment and diversification through the regular introduction of novel NA-HA pairings to different regions; and this creates significant challenges to the formulation of well-matched vaccines and other control efforts.

Viruses containing N2.1998 and N2.2002 lineage genes had multiple reassortment events, leading to the emergence of genetically distinct N2 clades. Our data demonstrated that N2 genes may persist at low detection levels, exchange HA genes via reassortment, and then through diversification and spatial dissemination the “minor” clades may rapidly become predominant. This is best exemplified by the frequency of detection of the N2.1998B and H1.Delta2 genes. The H1.Delta2 HA gene was originally paired with an N2.2002 when first detected in the US swine population, and this combination occurred at relatively low levels prior to 2011. In late 2011, a reassortment event that resulted in the pairing of the N2.1998B with the H1.Delta2 HA occurred, this coincided with correlated increases in the relative genetic diversity of both the H1.Delta2 and N2.1998B (Figure S4A), and subsequent substantive increases in the number of detections of the N2.1998B gene, with this combination representing 16% of H1 detections in 2017 (Zeller, Anderson, et al. 2018). Similarly, the N2.1998A clade persisted paired with H1.Delta1B subsequent to reassortment with H3.ClusterIVF, whereas H3.ClusterIVF IAV became infrequently detected. It is plausible to suggest that reassortment generating new NA-HA pairings could result in diversification of both HA and NA genes, adaptation, and potential wide dissemination. Our results suggested that NA-HA pairs and their evolution in swine are coordinated and likely play a role in alternating predominance of specific N2 clades and their paired HA clades.

An increase in detection frequency of specific NA-HA pairs following reassortment may be related to a favorable balance between the NA and HA protein activity (Hensley, et al. 2009), a better match between the surface genes and internal gene segments for replication (Gao, et al. 2017), or novel antigenic properties that evade population immunity (Abente, et al. 2016). The tanglegram in this study indicated that reassortment was followed by genetic divergence in connected NA and HA genes, evidenced in longer branch lengths and complete replacement of one NA-HA pairing with another over time. Newly paired HA and NA genes may require adaptation, and an additional collateral consequence of reassortment (antigenic shift) may be an increase in antigenic drift in both of the surface proteins. These data highlight the necessity to monitor genetic changes in both NA and HA, and patterns of NA and HA pairings through IAV surveillance in swine: these data will facilitate vaccine antigen selection and provide warning of potential rapid diversification that may result in a less effective vaccine formulation.

Our data revealed non-random pairing between NA and HA clades. We detected reassortment events that were a single (dead-end) detections or followed by multiple detections. The increased frequency of detection of novel NA-HA pairs following reassortment was indicative of sustained transmission of that NA-HA pairing within the swine population. Some reassortment events with subsequent transmission were also characterized by subsequent increased genetic diversity of both the N2 and HA in a coordinated manner based on the correlation in increase and decrease in relative genetic diversity across an 8 year period. A possible explanation for the coordinated changes in diversity genes is that the activity of these two proteins requires a balance for an IAV infection that results in transmission (Mitnaul, et al. 2000; Das, et al. 2011). The coordinated increase in diversity may arise due to selection pressure to optimize the HA and NA balance, evidence for this is a lag before the increase in detection numbers following reassortment, i.e., a novel NA and HA requires a period of adaptation (detected as an increase in relative genetic diversity) prior to increasing in detection frequency (Figure S3).

The genetic evolutionary rates of the N2 NA and the HA genes in swine were shown to have similar rates. In a limited number of studies, the mutation rate for the HA has been consistently similar to the NA across studies when measured empirically (Saitou and Nei 1986; Air, et al. 1990) and computationally (Khandaker, et al. 2013; Westgeest, et al. 2014). However, the general paradigm in IAV biology is a focus on the evolution of the HA and the role of immune-driven selection in its dynamics. Our data fit the growing body of literature that suggest that the NA evolves in a similar manner to the HA, and that immune driven selection may lead to functional amino acid changes (Correia, et al. 2018). We detected 26 amino acid sites that demonstrated a signal of diversifying selection, this included sites that were pervasively observed across different clades, some of which may have functional consequences. However, relative to data on selection analyses in HA genes, the number of sites in the NA undergoing diversifying selection is more limited, possibly as a compensatory mechanism for genetic changes in the HA.

The four monophyletic N2 clades circulating in U.S. swine had different geographic patterns and frequencies of reassortment. The N2 clades that were paired with more HA clades were the N2.2002A and N2.2002B clades, and these both had a higher relative genetic diversity when compared to the N2.1998A and B clades, as well as being frequently detected over the study period. This observation may reflect a “sampling effect” where a larger pool of genetic diversity in the N2 increases the likelihood of a reassortment event with an HA that may demonstrate onward transmission. Additionally, our data suggested that that genetic diversity increased following reassortment events; again, following reassortment, diversity may increase as a consequence of the genes being under coordinated evolutionary pressure to replicate within and transmit between hosts.

Detections of N2 clades were observed to have regional patterns. The N2.1998A was limited to 3 states in the Midwest, while the majority of N2.1998B detections were in the Southeast (North Carolina). The N2.2002A was detected across the Midwest, the N2.2002B was detected in the Midwest and Southeast. These patterns were derived from over eight years’ worth of data and support general swine agricultural practices where the Midwest appears to be a central point for pig shipment. Prior research suggested that the transmission of IAV in swine was observed in directed patterns in the US, namely from the Southwest and Southeast to the Midwest (Nelson, et al. 2011), and our data on the movement of the N2 genes is in general agreement with these prior findings. The implication of this is that directed movement of viruses could influence the opportunity of reassortment and subsequent patterns of diversity: areas that largely receive pigs and their viruses may provide more opportunities for reassortment. Consequently, a location such as Iowa would be expected to have the highest IAV genetic diversity as well as the most reassortment events. However, there are exceptions, some N2 clades remained geographically restricted despite interstate movement. The N2.1998B was introduced to the Midwest multiple times and to multiple states, but was not successfully established, indicating that factors (e.g., genetic compatibility or host-related) have prevented this clade from expanding across the US.

## Conclusion

This study demonstrates that changes within the NA and HA of the influenza virus in swine in the US were not independent of each other. While this study focused on the N2 due to its frequency of detection and apparent reassortment and genetic diversity, understanding the evolution of N1 and N2 subtypes will benefit vaccine development. If the HA vaccine antigen does not match currently circulating strains, a vaccine that contains a matched NA antigen may still reduce clinical signs and shedding of the virus. Further, monitoring the NA may serve as an early indicator of phenotypic change based on detected reassortment events. IAV reassortment may increase genetic and antigenic drift (Vijaykrishna, et al. 2011), and the current movement of swine between geographic locations in the US may increase these events. Future surveillance efforts with increased monitoring and surveillance of the genetic diversity of NA will be additive to the HA in control efforts to protect swine from influenza and reduce the chance for zoonotic transmission.

## Materials and Methods

### Data acquisition and phylogenetic clade assignment

Approximately 4000 paired nucleotide N2 NA and HA sequences detected in swine with accompanying metadata were downloaded from the Influenza Research Database (IRD) on October 18^th^, 2018 (Squires, et al. 2012; Yun Zhang, et al. 2017). Data was limited to IAV swine sequences from the US from August 9^th^, 2009 to September 20^th^, 2018. All NA sequences less than 1200 nt were removed (representing less that 85% of the coding region). Sequences associated with agricultural fair events and pigs were removed as they do not represent IAV from the general commercial swine population (i.e., sequences annotated with ‘OSU’) (Bowman, et al. 2017).

Phylogenetic clade classifications for H1 viruses were designated using the Swine H1 Clade Classification Tool provided on the IRD (Anderson, et al. 2016). H3 and NA clades were determined through maximum-likelihood phylogenetic analysis using reference sequences (Chang, et al. 2019). Reference nucleotide sequences and downloaded IAV from swine were aligned using MAFFT v7.27 (Katoh and Standley 2013). Maximum likelihood trees were inferred using FastTree2 v2.1.9 (Price, et al. 2010) for each gene alignment using a general time reversible model of nucleotide substitution with each IAV gene classified to clade based upon the nearest neighbor in the reference gene dataset. Within and between clade distances were calculated using MEGA X v10.1 (Kumar, et al. 2018).

### Quantifying relative genetic diversity in swine IAV N2 NA genes

Estimates of relative diversity for the N2.2002 and N2.1998 genetic lineages were determined through time-scaled Bayesian phylodynamic analyses and the inference of effective population size. Based upon computational limitations, we randomly sampled 620 genes from the N2.2002 lineage (from n=3650 total genes). These genes were aligned using MAFFT v7.27 (Katoh and Standley 2013) and a maximum-likelihood tree was inferred using FastTree2 v2.1.9 (Price, et al. 2010). Root-to-tip divergence was analyzed using TempEST v1.5.1 (Rambaut, et al. 2016) and genes with incongruent divergence and sampling date were removed (data available via GitHub). The resulting final dataset consisted of 600 non-identical N2.2002 genes. All of the available N2.1998 genes (n=596 taxa) were aligned, those that weren’t at least 50% of the gene were removed, and a maximum likelihood tree was inferred with root-to-tip divergence analyzed in TempEST v1.5.1 with two taxa removed as outliers, resulting in 583 taxa for analysis.

These datasets were analyzed using Bayesian phylogenetic methods in BEAST v1.8.4 (Drummond, et al. 2012) with the BEAGLE library v3.1.2 (Ayres, et al. 2011), implementing a generalized time reversible (GTR) nucleotide substitution model (Tavaré 1986) with gamma-distributed site heterogeneity (Yang 1994). We employed an uncorrelated relaxed clock with lognormal distribution (Drummond, et al. 2006), and a GMRF Bayesian skyride with time aware-smoothing as the coalescent model (Minin, et al. 2008). The MCMC chain length was set to 100 million iterations with sampling every 10,000 iterations. Results were analyzed using the GRMF skyride reconstruction in Tracer v1.6 (Rambaut and Drummond 2007).

### Measuring concurrent changes in the diversity of conserved NA and HA gene pairings

The frequency of particular NA and HA pairings was measured by splitting the data into the N2.2002 and N2.1998 monophyletic clades. NA-HA pairings that did not occur more than 10 times were removed from analysis, and the significance of gene pairings was measured with a post-hoc analysis following a Pearson’s chi square test for independence using the standard residuals in R v3.3.3 (R Core Team 2015). A confirmatory Fisher’s exact test was performed due to sparsity of some NA-HA pairings

Of these documented NA and HA pairings, N2.1998B was almost exclusively paired with H1.Delta2 HA genes (449 of 462 genes). Consequently, we analyzed the evolutionary dynamics of 462 taxa of N2.1998B clade NA genes and 522 taxa of H1.Delta2 HA genes separately. These genes were aligned, poor quality data were removed, and root-to-tip divergence was analyzed with divergent sequences removed. This process resulted in 458 N2.1998B genes and 517 H1.Delta2 genes (data available on GitHub). These data were analyzed using BEAST v1.8.4 implementing a GTR substitution model with gamma-distributed rate variation, an uncorrelated relaxed clock with lognormal distribution, and the GMRF Bayesian skyride coalescent. The MCMC chain was run for 100 million iterations with sampling every 10,000 iterations. The effective population size (EPS) for both clades N2.1998B and H1.Delta2 was plotted to determine correlated changes in relative genetic diversity. A similar analysis with the same analytical settings was conducted on the N2.2002A genes that were paired with H1.Delta1B HA (following screening, n=842 N2.2002A and H1.Delta1B taxa for analysis).

### Estimating the evolutionary rate of swine IAV N2 NA and HA genes

To estimate the evolutionary rate of N2 NA, we created alignments of each of the major of genetic clades: N2.1998A, N2.1998B, N2.2002A, and N2.2002B (n = 115, 458, 262, 310, respectively). To estimate the evolutionary rate of H1 and H3 HA, we generated alignments of H1.Gamma, H1.Delta2, H3.2010.1, and H3.ClusterIVA (n=300, 300, 299, 298). Using BEAST 2 v2.5.1 (Bouckaert, et al. 2014), we implemented an unlinked codon substitution model split into independent partitions (1 + 2 + 3) and unlinked clock models for each codon position. The analysis used a general time reversible (GTR) nucleotide substitution model with a gamma site heterogeneity, an uncorrelated relaxed clock with lognormal distribution, and an exponential population growth coalescent tree prior. The MCMC chain length was set to 100 million iterations with sampling every 10,000 iterations. Mean rate of substitution was calculated by averaging the values for the three codon position’s mean rate of substitution in terms of sites/mutation/year with distribution for each and visualized using ggplot2 (Wickham 2016).

### Detecting diversification following reassortment of N2 NA in swine IAV

Reassortment in N2 NA gene was identified using topological incongruity in gene trees (Boni, et al. 2010). Maximum likelihood trees were inferred for N2.1998 and N2.2002 using RAxML v8.2.11 (Stamatakis 2014) with a GTR model with a gamma-distributed rate variation and statistical support values determined using rapid bootstrapping with the autoMRE criterion. Trees were visualized in FigTree v1.4.3 (Rambaut 2012), and we defined sustained transmission of a reassorted pairing as one where a monophyletic clade of NA genes acquired a novel HA genetic clade, and the NA-HA combination was maintained within the monophyletic clade for more than 10 subsequent branches. To facilitate visualization, a randomly sampled set of 300 HA and NA genes were sorted and transformed into proportional branch length cladograms and built into a tanglegram in R v3.3.3 (R Core Team 2015) using the dendextend v1.9.0 package (Galili 2015). Branches on these constituent maximum-likelihood trees were colored by phylogenetic clades with lines connecting trees colored based on HA clade to emphasize reassortment events.

Branch lengths of the NA tree were compared to assess whether reassortment affected diversity. To achieve this, we identified and extracted monophyletic clades that contained a reassortment event. Extracted clades were realigned using MAFFT v7.27 (Katoh and Standley 2013), and a maximum likelihood trees were inferred using FastTree2 v2.1.9 (Price, et al. 2010). We were able to generate 7 phylogenies that demonstrated an NA gene acquiring a novel HA gene that then persisted in the US swine population. The 7 trees were loaded into R v3.3.3, and the patristic distance from each NA gene to the oldest leaf node was determined using APE v5.2 (Paradis, et al. 2004). A linear model of y = β0 + β1x was fitted to these data, where y was patristic distance, β1 was date of collection of the NA gene, and β0 represents an intercept. A linear model was fit for patristic distance for NA genes that had the prior paired HA clade, and for the NA genes that had acquired a novel HA gene. The fitted models for each of reassortment event (i.e., 14 models, 2 for each of the 7 reassortment events) were compared using the Chow Test from the gap package v1.2.2 in R (Zhao 2007). To determine whether using one or two linear models was appropriate, we measured whether there was significant reduction in the residuals at the 95% confidence level.

Within the NA genes, we used MEME (Murrell, et al. 2012), part of the HyPhy package v2.3.14, to test for instances of episodic and pervasive selection by using a mixed-effects model of evolution. Results from MEME were confirmed by conducting concurrent analyses with by FEL and SLAC (Kosakovsky Pond and Frost 2005), and reviewed via HyPhy vision v2.4.3.

### Inferring the directionality of N2 and HA genes following reassortment in swine IAV

To quantitatively characterize the direction of the exchange of NA and HA genes, we adapted a phylogeographic approach, using NA genetic clade information as our “location state.” The same data subset and setup was used as in our analyses on quantifying changes in relative genetic diversity. Each NA gene had an HA genetic clade designated as a trait, and transitions from one category to another (e.g., from N2.1998B/H1.Gamma to N2.1998B/H3 2010.1) were inferred along the internal branches representing the evolutionary history of the virus. These transitions are termed Markov jumps and state change counts for the HA clade trait were reconstructed in BEAST v1.8.4. The sites for the trait were calculated using an asymmetric substitution model, a Bayesian stochastic search variable selection (BSSVS) (Edwards, et al.), and an uncorrelated relaxed clock model with lognormal distribution. Analysis output was assessed in Tracer v1.6.0 to confirm convergence and SPREAD3 v0.9.6 (Bielejec, et al. 2016) was used to compute the Bayes factors with 10% burn-in. Bayes factors were plotted using the inferno color scheme from the R viridis package v0.5.1 using the ggplot2 v3.1.0 and cowplot v0.9.4.

### Assessing the spatial distribution of N2 NA genes in swine IAV

Swine IAV N2 data were reduced to subsets that included only sequences that included a US state. The number of detections of each N2 genetic clade was rendered on a US map using the R maps module v3.3.0, colored to emphasize differences using the viridis color scheme and rendered by ggplot2 v3.1.0. Statistical association between N2 genetic clade and state of collection was assessed with a Pearson’s chi square test for independence, and then a post hoc analysis of chi square test using the standard residuals in R v3.3.3 (R Core Team 2015). A Fisher’s exact test was performed to confirm the results of the initial chi square test. To quantify the movement of the NA genes across the US, we used a phylogeographic approach with US state as our “location state.” Each NA gene had an US state designated as a trait, and transitions from one category to another (e.g., from North Carolina to Iowa) were inferred along the internal branches representing the evolutionary history of the virus. These number of state change counts for the US state trait were reconstructed in BEAST v1.8.4 following the same methods described above.

## Acknowledgements

This work was supported by the Iowa State University Presidential Interdisciplinary Research Initiative; the Iowa State University Veterinary Diagnostic Laboratory; the U.S. Department of Agriculture (USDA) Agricultural Research Service (ARS project number 5030-32000-120-00-D); an National Institute of Allergy and Infectious Diseases (NIAID) at the National Institutes of Health interagency agreement associated with the Center of Research in Influenza Pathogenesis, an NIAID funded Center of Excellence in Influenza Research and Surveillance (grant HHSN272201400008C to A.L.V.); the USDA Agricultural Research Service Research Participation Program of the Oak Ridge Institute for Science and Education (ORISE) through an interagency agreement between the U.S. Department of Energy (DOE) and USDA Agricultural Research Service (contract number DE-AC05-06OR23100 to J.C.); and the SCINet project of the USDA Agricultural Research Service (ARS project number 0500-00093-001-00-D). The funders had no role in study design, data collection and interpretation, or the decision to submit the work for publication. Mention of trade names or commercial products in this article is solely for the purpose of providing specific information and does not imply recommendation or endorsement by the USDA, DOE, or ORISE. USDA is an equal opportunity provider and employer.

## Supplementary Figure Legends

**Figure S1**. Effective population size representing the relative genetic diversity of the N2 clades detected in North American swine from 2009-2018, N2.1998A, N2.1998B, N2.2002A, and N2.2002B. Temporal dominance in genetic diversity of NA N2 was apparent after separating the clades. The N2.2002A genetic diversity peaked in mid-2012, N2.2002B peaked in mid-2015, and N2.1998B peaked at approximately mid-2016. N2.2002A and N2.2002.B maintained moderate genetic diversity at the end of the study period.

**Figure S2.** Regression models fitted to patristic distance from the oldest gene on a phylogenetic tree for neuraminidase (NA) from a donor hemagglutinin in red (HA) to a recipient HA in blue. The slope of the regression line is representative of the mutation rate of the NA paired with either the donor or recipient HA. (A) IAV containing N2.1998A paired with H3.ClusterIVF donating NA genes to IAV with H1.Delta1B (Figure 2, reassortment event 1), (B) N2.2002A paired with H1.Delta1B donating NA genes to H3.2010.1 (Figure 2, reassortment event 4), (C) N2.2002A paired with H3.ClusterIVB donating NA genes to IAV with H1.Alpha (Figure 2, reassortment event 5), (D) N2.2002A paired with H1.Alpha donating NA genes to IAV with H3.2010.1 (Figure 2, reassortment event 6), (E) N2.2002B paired with H3.ClusterIVB donating NA genes to IAV with H3.ClusterIVA (Figure 2, reassortment event 7), (F) N2.2002B paired with H3.ClusterIVA donating NA genes to IAV with H1.Delta1A (Figure 2, reassortment event 8), (G) N2.2002B paired with H3.ClusterIVA donating NA genes to IAV with H1.2010.2 (Figure 2, reassortment event 9). The Chow test indicated a significant break, indicating that two linear models was more valid than one for subsets A, E, and F. Linear breaks may indicate a temporal shift in mutation rate.

**Figure S3.** Neuraminidase clades over time paired with (A) H1.Delta1a, (B) H3.ClusterIVA, (C) H3.2010.1 in IAV detected from 2009-2018. Graphs depict the number of detections of each HA and NA pairing. A change in predominant HA and NA pairings occurred over time.

**Figure S4.** (A) The observed number of HA and N2 neuraminidase clade pairings represented in the dataset. (B) The standardized residuals of the HA and N2 neuraminidase clade pairings based on the chi squared test for independence (p < 0.001) performed separately on the N2.1998 and N2.2002 clades. The post hoc analysis demonstrated individual contribution to the p-value of each clade pairing. Red cells represent HA-N2 pairing observed more than expected and blue cells represent pairings observed less than expected if pairings were random. The results of this analysis support that HA pairing with N2 was not random.

**Figure S5.** (A) Relative diversity of the N2.1998B clade and the H1.Delta2 clade over time from 2009-2018. Relative diversity of H1.Delta2 and N2.1998B was not correlated until after 2012, prior to which the H1.Delta2 was primarily paired with N2.2002. (B) Relative diversity of the N2.2002A clade and the H1.Delta1B clade that were paired from 2009-2018. Median EPS were denoted by lines with the 95% higher posterior density shaded in the same color. Temporally matched changes at similar magnitudes suggested a correlation between the diversity of the shown NA-HA.

**Figure S6.** The mean nucleotide substitution rate of H1, H3, and N2 clades. Substitution rates were calculated using BEAST2. Each codon position represented a different log normal relaxed clock model, and a GTR+ Γ substitution model with four categories. The three clock rates for each codon position were averaged to determine the mean nucleotide substitution rate across the entire gene. N2.2002A had the lowest mean substitution rate at 0.0041, while H3.2010.1 had the highest substitution rate at 0.0050. The mean HA substitution rate was 0.0005 nucleotides/site/year faster than the NA mean substitution rate.

## References

Abente EJ, Santos J, Lewis NS, Gauger PC, Stratton J, Skepner E, Anderson TK, Rajao DS, Perez DR, Vincent AL. 2016. The molecular determinants of antibody recognition and antigenic drift in the H3 hemagglutinin of swine influenza A virus. Journal of virology 90:8266–8280.

Air GM, Gibbs AJ, Laver WG, Webster RG. 1990. Evolutionary changes in influenza B are not primarily governed by antibody selection. Proceedings of the National Academy of Sciences 87:3884–3888.

Anderson TK, Macken CA, Lewis NS, Scheuermann RH, Van Reeth K, Brown IH, Swenson SL, Simon G, Saito T, Berhane Y. 2016. A Phylogeny-Based Global Nomenclature System and Automated Annotation Tool for H1 Hemagglutinin Genes from Swine Influenza A Viruses. mSphere 1:e00275–00216.

Anderson TK, Nelson MI, Kitikoon P, Swenson SL, Korslund JA, Vincent AL. 2013. Population dynamics of cocirculating swine influenza A viruses in the United States from 2009 to 2012. Influenza Other Respir Viruses 7 Suppl 4:42–51.

Ayres DL, Darling A, Zwickl DJ, Beerli P, Holder MT, Lewis PO, Huelsenbeck JP, Ronquist F, Swofford DL, Cummings MP. 2011. BEAGLE: an application programming interface and high-performance computing library for statistical phylogenetics. Systematic biology 61:170–173.

Bielejec F, Baele G, Vrancken B, Suchard MA, Rambaut A, Lemey P. 2016. SpreaD3: interactive visualization of spatiotemporal history and trait evolutionary processes. Molecular biology and evolution 33:2167–2169.

Boni MF, de Jong MD, van Doorn HR, Holmes EC. 2010. Guidelines for identifying homologous recombination events in influenza A virus. PloS one 5.

Bouckaert R, Heled J, Kühnert D, Vaughan T, Wu C-H, Xie D, Suchard MA, Rambaut A, Drummond AJ. 2014. BEAST 2: a software platform for Bayesian evolutionary analysis. PLoS computational biology 10:e1003537.

Bowman AS, Walia RR, Nolting JM, Vincent AL, Killian ML, Zentkovich MM, Lorbach JN, Lauterbach SE, Anderson TK, Davis TC. 2017. Influenza A (H3N2) virus in swine at agricultural fairs and transmission to humans, Michigan and Ohio, USA, 2016. Emerging infectious diseases 23:1551.

Chang J, Anderson TK, Zeller MA, Gauger PC, Vincent AL. 2019. octoFLU: Automated Classification for the Evolutionary Origin of Influenza A Virus Gene Sequences Detected in US Swine. Microbiology resource announcements 8:e00673–00619.

Chow GC. 1960. Tests of equality between sets of coefficients in two linear regressions. Econometrica: Journal of the Econometric Society:591–605.

Correia V, Abecasis AB, Rebelo-de-Andrade H. 2018. Molecular footprints of selective pressure in the neuraminidase gene of currently circulating human influenza subtypes and lineages. Virology 522:122–130.

Das SR, Hensley SE, David A, Schmidt L, Gibbs JS, Puigbò P, Ince WL, Bennink JR, Yewdell JW. 2011. Fitness costs limit influenza A virus hemagglutinin glycosylation as an immune evasion strategy. Proceedings of the National Academy of Sciences 108:E1417–E1422.

Dawood FS. 2009. Emergence of a Novel Swine-Origin Influenza A (H1N1) Virus in Humans. The New England Journal of Medicine 360:2605–2615.

Drummond AJ, Ho SY, Phillips MJ, Rambaut A. 2006. Relaxed phylogenetics and dating with confidence. PLoS biology 4:e88.

Drummond AJ, Suchard MA, Xie D, Rambaut A. 2012. Bayesian phylogenetics with BEAUti and the BEAST 1.7. Molecular biology and evolution 29:1969–1973.

Dykhuis-Haden C, Painter T, Fangman T, Holtkamp D. 2012. Assessing production parameters and economic impact of swine influenza, PRRS and *Mycoplasma hyopneumoniae* on finishing pigs in a large production system. American Association of Swine Veterinarians; Denver, Colorado. p. 75–76.

Edwards CJ, Suchard MA, Lemey P, Welch JJ, Barnes I, Fulton TL, Barnett R, O’Connell TC, Coxon P, Monaghan N. Supplemental Information Ancient Hybridization and an Irish Origin for the Modern Polar Bear Matriline.

Galili T. 2015. dendextend: an R package for visualizing, adjusting and comparing trees of hierarchical clustering. Bioinformatics 31:3718–3720.

Gao S, Anderson TK, Walia RR, Dorman KS, Janas-Martindale A, Vincent AL. 2017. The genomic evolution of H1 influenza A viruses from swine detected in the United States between 2009 and 2016. Journal of General Virology 98:2001–2010.

Gramer M. 2006. Swine influenza virus: the only constant is change.

Hensley SE, Das SR, Bailey AL, Schmidt LM, Hickman HD, Jayaraman A, Viswanathan K, Raman R, Sasisekharan R, Bennink JR. 2009. Hemagglutinin receptor binding avidity drives influenza A virus antigenic drift. Science 326:734–736.

Katoh K, Standley DM. 2013. MAFFT multiple sequence alignment software version 7: improvements in performance and usability. Molecular biology and evolution 30:772–780.

Khandaker I, Suzuki A, Kamigaki T, Tohma K, Odagiri T, Okada T, Ohno A, Otani K, Sawayama R, Kawamura K. 2013. Molecular evolution of the hemagglutinin and neuraminidase genes of pandemic (H1N1) 2009 influenza viruses in Sendai, Japan, during 2009–2011. Virus Genes 47:456–466.

Koen J. 1919. A practical method for field diagnosis of swine diseases. Am J Vet Med 14:468–470.

Kosakovsky Pond SL, Frost SD. 2005. Not so different after all: a comparison of methods for detecting amino acid sites under selection. Molecular biology and evolution 22:1208–1222.

Kumar S, Stecher G, Li M, Knyaz C, Tamura K. 2018. MEGA X: molecular evolutionary genetics analysis across computing platforms. Molecular biology and evolution 35:1547–1549.

Minin VN, Bloomquist EW, Suchard MA. 2008. Smooth skyride through a rough skyline: Bayesian coalescent-based inference of population dynamics. Molecular biology and evolution 25:1459–1471.

Mitnaul LJ, Matrosovich MN, Castrucci MR, Tuzikov AB, Bovin NV, Kobasa D, Kawaoka Y. 2000. Balanced hemagglutinin and neuraminidase activities are critical for efficient replication of influenza A virus. Journal of virology 74:6015–6020.

Monto AS, Kendal AP. 1973. Effect of neuraminidase antibody on Hong Kong influenza. The Lancet 301:623–625.

Murrell B, Wertheim JO, Moola S, Weighill T, Scheffler K, Kosakovsky Pond SL. 2012. Detecting individual sites subject to episodic diversifying selection. PLoS Genet 8:e1002764.

Nelson MI, Lemey P, Tan Y, Vincent A, Lam TT-Y, Detmer S, Viboud C, Suchard MA, Rambaut A, Holmes EC. 2011. Spatial dynamics of human-origin H1 influenza A virus in North American swine. PLoS pathogens 7:e1002077.

Neverov AD, Lezhnina KV, Kondrashov AS, Bazykin GAJPg. 2014. Intrasubtype reassortments cause adaptive amino acid replacements in H3N2 influenza genes. 10:e1004037.

Paradis E, Claude J, Strimmer K. 2004. APE: analyses of phylogenetics and evolution in R language. Bioinformatics 20:289–290.

Price MN, Dehal PS, Arkin AP. 2010. FastTree 2–approximately maximum-likelihood trees for large alignments. PLoS One 5:e9490.

R Core Team. 2015. R: A language and environment for statistical computing.

Rambaut A. 2012. FigTree v1. 4. Molecular evolution, phylogenetics and epidemiology. Edinburgh, UK: University of Edinburgh, Institute of Evolutionary Biology.

Rambaut A, Drummond AJ. 2007. Tracer v1. 6 http://beast. bio. ed. ac. uk. In: Tracer.

Rambaut A, Lam TT, Max Carvalho L, Pybus OG. 2016. Exploring the temporal structure of heterochronous sequences using TempEst (formerly Path-O-Gen). Virus evolution 2:vew007.

Saitou N, Nei M. 1986. Polymorphism and evolution of influenza A virus genes. Molecular biology and evolution 3:57–74.

Sandbulte MR, Gauger PC, Kitikoon P, Chen H, Perez DR, Roth JA, Vincent AL. 2016. Neuraminidase inhibiting antibody responses in pigs differ between influenza A virus N2 lineages and by vaccine type. Vaccine 34:3773–3779.

Sandbulte MR, Spickler AR, Zaabel PK, Roth JA. 2015. Optimal Use of Vaccines for Control of Influenza A Virus in Swine. Vaccines (Basel) 3:22–73.

Smith GJ, Vijaykrishna D, Bahl J, Lycett SJ, Worobey M, Pybus OG, Ma SK, Cheung CL, Raghwani J, Bhatt S, et al. (1709 co-authors). 2009. Origins and evolutionary genomics of the 2009 swine-origin H1N1 influenza A epidemics. Nature 459:1122–1125.

Squires RB, Noronha J, Hunt V, Garcia-Sastre A, Macken C, Baumgarth N, Suarez D, Pickett BE, Zhang Y, Larsen CN, et al. 2012. Influenza research database: an integrated bioinformatics resource for influenza research and surveillance. Influenza Other Respir Viruses 6:404–416.

Stamatakis A. 2014. RAxML version 8: a tool for phylogenetic analysis and post-analysis of large phylogenies. Bioinformatics 30:1312–1313.

Tavaré S. 1986. Some probabilistic and statistical problems in the analysis of DNA sequences. Lectures on mathematics in the life sciences 17:57–86.

Vijaykrishna D, Smith GJ, Pybus OG, Zhu H, Bhatt S, Poon LL, Riley S, Bahl J, Ma SK, Cheung CL. 2011. Long-term evolution and transmission dynamics of swine influenza A virus. Nature 473:519.

Vincent AL, Lager KM, Janke BH, Gramer MR, Richt JA. 2008. Failure of protection and enhanced pneumonia with a US H1N2 swine influenza virus in pigs vaccinated with an inactivated classical swine H1N1 vaccine. Veterinary microbiology 126:310–323.

Vincent AL, Ma W, Lager KM, Gramer MR, Richt JA, Janke BH. 2009. Characterization of a newly emerged genetic cluster of H1N1 and H1N2 swine influenza virus in the United States. Virus Genes 39:176–185.

Vincent AL, Ma W, Lager KM, Janke BH, Richt JA. 2008. Swine influenza viruses: a North American perspective. Advances in virus research 72:127–154.

Vincent AL, Perez DR, Rajao D, Anderson TK, Abente EJ, Walia RR, Lewis NS. 2017. Influenza A virus vaccines for swine. Vet Microbiol 206:35–44.

Wang Y, Gagnon CA, Savard C, Music N, Srednik M, Segura M, Lachance C, Bellehumeur C, Gottschalk M. 2013. Capsular sialic acid of Streptococcus suis serotype 2 binds to swine influenza virus and enhances bacterial interactions with virus-infected tracheal epithelial cells. Infection and immunity 81:4498–4508.

Westgeest KB, Russell CA, Lin X, Spronken MI, Bestebroer TM, Bahl J, van Beek R, Skepner E, Halpin RA, de Jong JC. 2014. Genomewide analysis of reassortment and evolution of human influenza A (H3N2) viruses circulating between 1968 and 2011. Journal of virology 88:2844–2857.

Wickham H. 2016. ggplot2: elegant graphics for data analysis: Springer.

Yang Z. 1994. Maximum likelihood phylogenetic estimation from DNA sequences with variable rates over sites: approximate methods. Journal of Molecular evolution 39:306–314.

Yun Zhang, Brian D. Aevermann, Tavis K. Anderson, David F. Burke, Gwenaelle Dauphin, Zhiping Gu, Sherry He, Sanjeev Kumar, Christopher N. Larsen, Alexandra J. Lee, et al. 2017. Influenza Research Database: An integrated bioinformatics resource for influenza virus research. Nucleic Acids Res 45:D466–D474.

Zeller MA, Anderson TK, Walia RW, Vincent AL, Gauger PC. 2018. ISU FLU ture: a veterinary diagnostic laboratory web-based platform to monitor the temporal genetic patterns of Influenza A virus in swine. BMC Bioinformatics 19:397.

Zeller MA, Li G, Harmon KM, Zhang J, Vincent AL, Anderson TK, Gauger PC. 2018. Complete Genome Sequences of Two Novel Human-Like H3N2 Influenza A Viruses, A/swine/Oklahoma/65980/2017 (H3N2) and A/Swine/Oklahoma/65260/2017 (H3N2), Detected in Swine in the United States. Microbiol Resour Announc 7:e01203–01218.

Zhao JH. 2007. A Genetic Analysis Package with R. Journal of Statistical Software 23:1–18.

Zhou NN, Senne DA, Landgraf JS, Swenson SL, Erickson G, Rossow K, Liu L, Yoon K-j, Krauss S, Webster RG. 1999. Genetic reassortment of avian, swine, and human influenza A viruses in American pigs. Journal of virology 73:8851–8856.

