## Supplemental Figures S1-S6 for "Coordinated evolution between N2 neuraminidase and H1 and H3 hemagglutinin genes increased influenza A virus genetic diversity in swine"

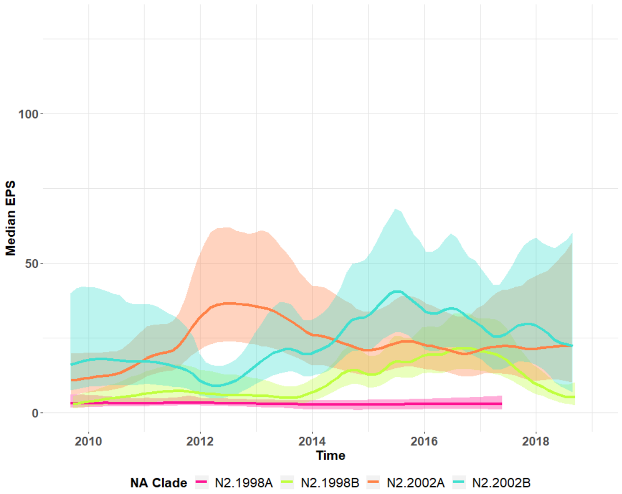


Figure S1. Effective population size representing the relative genetic diversity of the N2 clades detected in North American swine from 2009-2018, N2.1998A, N2.1998B, N2.2002A, and N2.2002B. Temporal dominance in genetic diversity of NA N2 was apparent after separating the clades. The N2.2002A genetic diversity peaked in mid-2012, N2.2002B peaked in mid-2015, and N2.1998B peaked at approximately mid-2016. N2.2002A and N2.2002.B maintained moderate genetic diversity at the end of the study period.

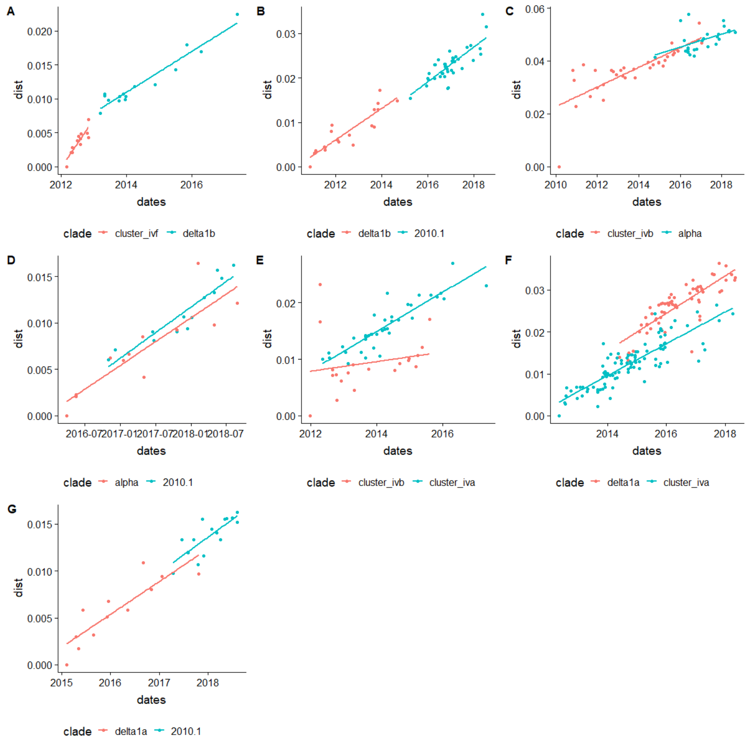


Figure S2.Regression models fitted to patristic distance from the oldest gene on a phylogenetic tree for neuraminidase (NA) from a donor hemagglutinin in red (HA) to a recipient HA in blue. The slope of the regression line is representative of the mutation rate of the NA paired with either the donor or recipient HA. (A) IAV containing N2.1998A paired with H3.ClusterIVF donating NA genes to IAV with H1.Delta1B (Figure 2, reassortment event 1), (B) N2.2002A paired with H1.Delta1B donating NA genes to H3.2010.1 (Figure 2, reassortment event 4), (C) N2.2002A paired with H3.ClusterIVB donating NA genes to IAV with H1.Alpha (Figure 2, reassortment event 5), (D) N2.2002A paired with H1.Alpha donating NA genes to IAV with H3.2010.1 (Figure 2, reassortment event 6), (E) N2.2002B paired with H3.ClusterIVB donating NA genes to IAV with H3.ClusterIVA (Figure 2, reassortment event 7), (F) N2.2002B paired with H3.ClusterIVA donating NA genes to IAV with H1.Delta1A (Figure 2, reassortment event 8), (G) N2.2002B paired with H3.ClusterIVA donating NA genes to IAV with H1.2010.2 (Figure 2, reassortment event 9). The Chow test indicated a significant break, indicating that two linear models was more valid than one for subsets A, E, and F. Linear breaks may indicate a temporal shift in mutation rate.

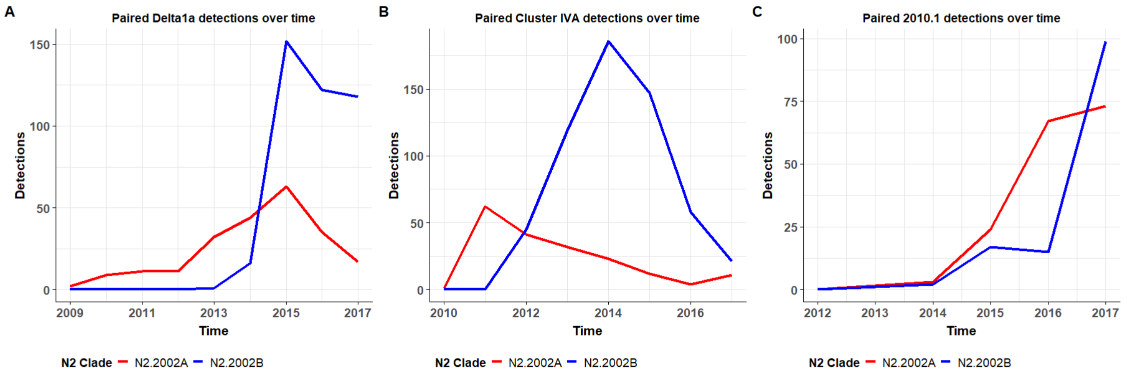


Figure S3.Neuraminidase clades over time paired with (A) H1.Delta1a, (B) H3.ClusterIVA, (C) H3.2010.1 in IAV detected from 2009-2018. Graphs depict the number of detections of each HA and NA pairing. A change in predominant HA and NA pairings occurred over time.

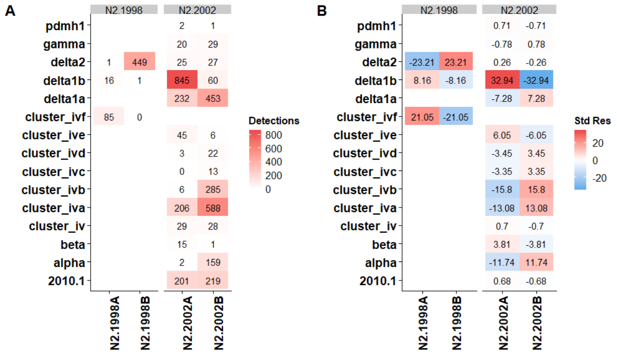


Figure S4. (A) The observed number of HA and N2 neuraminidase clade pairings represented in the dataset. (B) The standardized residuals of the HA and N2 neuraminidase clade pairings based on the chi squared test for independence ($p < 0.001$) performed separately on the N2.1998 and N2.2002 clades. The post hoc analysis demonstrated individual contribution to the p-value of each clade pairing. Red cells represent HA-N2 pairing observed more than expected and blue cells represent pairings observed less than expected if pairings were random. The results of this analysis support that HA pairing with N2 was not random.

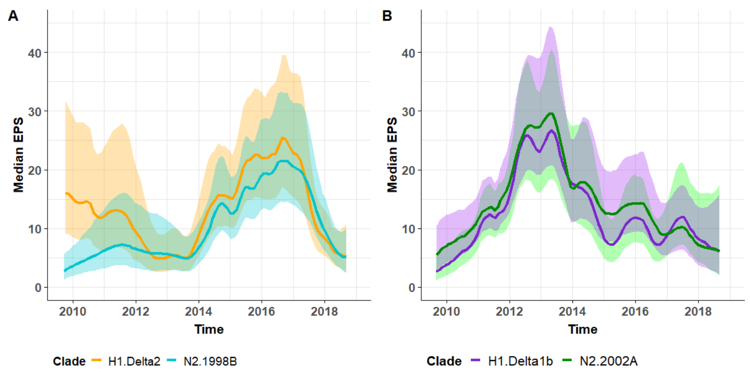


Figure S5. (A) Relative diversity of the N2.1998B clade and the H1.Delta2 clade over time from 2009-2018. Relative diversity of H1.Delta2 and N2.1998B was not correlated until after 2012, prior to which the H1.Delta2 was primarily paired with N2.2002 . (B) Relative diversity of the N2.2002A clade and the H1.Delta1B clade that were paired from 2009-2018. Median EPS were denoted by lines with the 95% higher posterior density shaded in the same color. Temporally matched changes at similar magnitudes suggested a correlation between the diversity of the shown NA-HA.

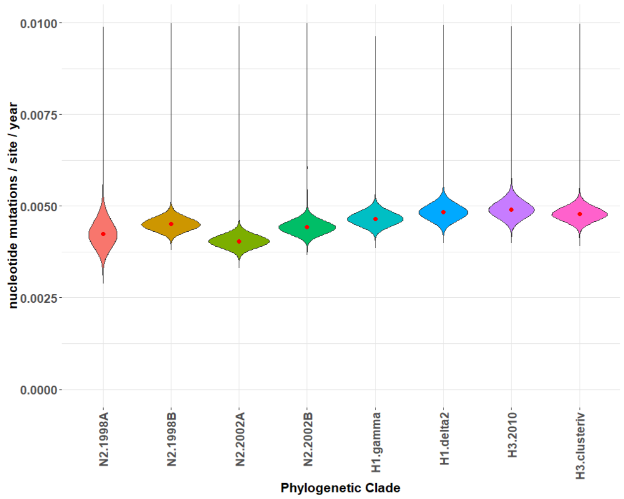


Figure S6. The mean nucleotide substitution rate of H1, H3, and N2 clades. Substitution rates were calculated using BEAST2. Each codon position represented a different log normal relaxed clock model, and a GTR+ Γ substitution model with four categories. The three clock rates for each codon position were averaged to determine the mean nucleotide substitution rate across the entire gene. N2.2002A had the lowest mean substitution rate at 0.0041, while H3.2010.1 had the highest substitution rate at 0.0050. The mean HA substitution rate was 0.0005 nucleotides/site/year faster than the NA mean substitution rate.
